# Origins and Global Context of *Brucella abortus* in Italy

**DOI:** 10.1101/077529

**Authors:** Giuliano Garofolo, Elisabetta Di Giannatale, Ilenia Platone, Katiuscia Zilli, Lorena Sacchini, Anna Abass, Massimo Ancora, Cesare Cammà, Guido Di Donato, Fabrizio De Massis, Paolo Calistri, Kevin P. Drees, Jeffrey T. Foster

**Author notes:** Cesare Cammà.

## Abstract

**Background:** Brucellosis is a common and chronic disease of cattle and other bovids that often causes reproductive disorders. Natural infection in cattle is caused by *Brucella abortus* and transmission typically occurs during abortions, calving, or nursing. Brucellosis is also a major zoonotic disease due to contamination of dairy products or contact with the tissues of infected animals. Brucellosis has been eradicated from most of the developed world in the last forty years but persists in many regions; *B. abortus* remains prevalent in portions of Africa, the Middle East, Asia, and Central and South America, as well as in the Mediterranean basin. Specifically, *B. abortus* has persisted in southern Italy in both cattle and water buffalo. Previous attempts at analyzing the genetic structure of *B. abortus* in Italy have been challenging due to the limited genetic variability and unresolved global population structure of this pathogen.

**Results:** We conducted genome-wide phylogenetic analysis on 11 representative strains of *B. abortus* from Italy, and compared these sequences to a worldwide collection of publically available genomes. Italian isolates belong to three clades basal to the global *B. abortus* lineage. Using six assays designed to identify substructure within the Italian clades in a collection of 261 isolates, one clade predominates throughout endemic districts in the country, while the other two clades are more geographically restricted to southern Italy.

**Conclusions:** Although related strains exist worldwide, *B. abortus* isolates from Italy are substantially different than those found in much of the rest of Europe and North America, and are more closely related to strains from the Middle East and Asia. Our assays targeting genetic substructure within Italy allowed us to identify the major lineages quickly and inexpensively, without having to generate whole genome sequences for a large isolate collection. These findings highlight the importance of genetic studies to assess the status and the history of pathogens.

## Introduction

Although eradicated throughout much of the developed world, bovine brucellosis continues to be common in Southern Italy [1]. *Brucella abortus,* the etiological agent, is a Gram-negative non-motile and non-sporulating bacterium that forms coccobacilli with oxidative metabolism [2]. At the beginning of the 20th century, *B. abortus* was recognized by Benhard Bang as the causative agent of epizootic abortion in cattle, but only later was this disease associated with Malta fever of goats caused by *B. melitensis* [3] and classified by Meyer in the genus *Brucella* [4]. The disease presents primarily as reproductive disorders in cattle, resulting in substantial economic losses to agriculture [5]. For example, the economic burden of bovine brucellosis in Latin America has been estimated at $600 million annually [6]. *Brucella abortus* is a significant zoonotic agent, and humans are typically infected by consumption of raw dairy products, at slaughter, or during veterinary care and animal husbandry [7].

The Italian government has conducted a country-wide eradication program for cattle brucellosis since 1994 [8]. Animal cases are currently limited to seven regions of southern Italy, with the highest prevalence of infection in areas of Sicily, Calabria, and Apulia regions [9]. This eradication program, along with strict regulations on cattle movements, has reduced the prevalence and geographical distribution of brucellosis in Italy [10]. In order to achieve the eradication of brucellosis, the identification of the sources of infection is essential, as is determining whether the same strains are circulating within a region or continuously reintroduced from other areas or countries. Additionally, the host species and source region are important for the epidemiology of human zoonoses.

Molecular epidemiology has been challenging in *Brucella* due to relatively high levels of nucleotide similarity throughout the genus [11]. Traditional molecular approaches involving biochemical testing are only able to categorize samples into broad phenotypic groups of biovars, and for *B. abortus* eight different biovars have been described (biovars 1–7 and 9) [12]. More recent genetic analyses using variable number tandem repeat (VNTR) markers have given much higher resolution and can be highly informative in some cases [13–16], but homoplasy in VNTR markers and a lack of sampling for some regions make it challenging to place *B. abortus* into a true global phylogeny [17].

We used whole genome sequencing (WGS) to place 11 Italian *B. abortus* isolates onto a global context. We also identified specific single nucleotide polymorphisms (SNPs) that define the three major Italian clades in our study, and then developed clade-specific assays to genotype our Italian collection of 261 isolates. This genotyping allowed us to rapidly and inexpensively identify the lineages of *B. abortus* circulating in southern Italy and place Italian strains in a global phylogeny. In Italy, where brucellosis is a major public health concern, the information retrieved by this genotyping will also support the epidemiological investigations in case of human infections.

## Results

Whole genome comparisons generated approximately 9,000 putative SNPs that were used to construct a maximum parsimony phylogenetic tree. Italian isolates of *B. abortus* were part of three clades that are basal to the most common worldwide lineage, which we refer to as the biovar 1, 2, 4 lineage (hereafter biovar 1/2/4), that is ubiquitous in North America and parts of Europe (Fig. 1). Isolates from western Italy and a closely related sample from France form a “West Italia” subclade defined by 83 SNPs. A broader clade that also includes isolates from Poland, Spain, and China is defined by 91 SNPs. Members of the West Italia subclade and broader clade are closely related to the biovar 1/2/4 lineage, which is consistent with the biochemical designation of most of these isolates as biovar 1.

**Fig. 1.**
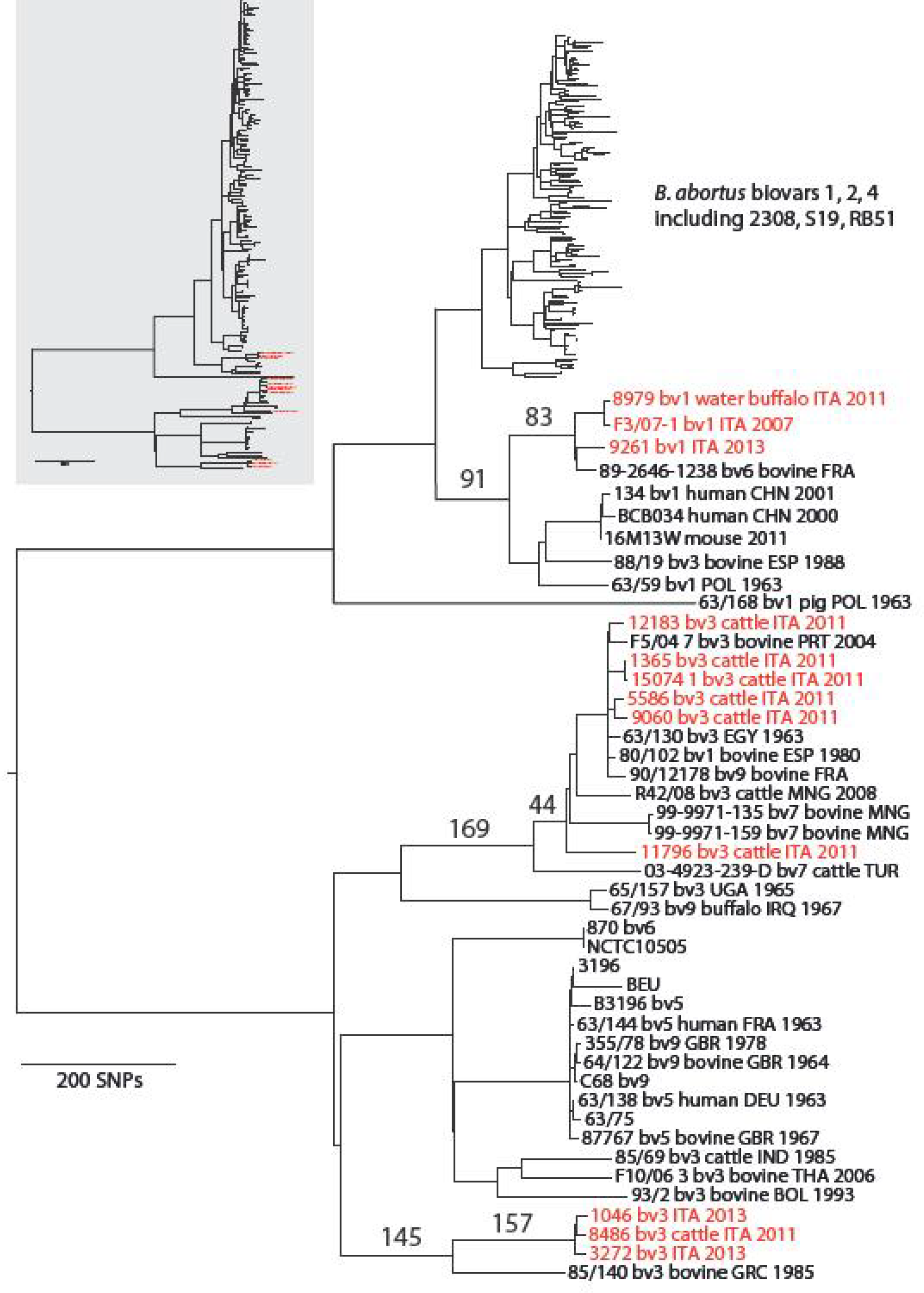
Maximum parsimony tree for *B. abortus,* with genomes from Italian isolates highlighted in red. Bootstrap support values were 100 for all major branches, with only the shallowest clades having less support (data not shown). Numbered branches indicate the number of SNPs identified for the clade (left most) and subclade (right most) level groupings for the West Italia (top), Trans Italia (middle) and East Italia (bottom) subclades. Samples are identified by the strain name and where known, the biovar (bv) number, animal source, country of origin (3 letter ISO codes), and year of isolation.

The other Italian isolates are part of an even more basal and considerably more diverse group of *B. abortus* genomes. Unlike the biovar 1/2/4 lineage, this basal group is comprised largely of isolates from biovars 3, 5, 6, 7, 9 that are broadly distributed across Europe, Asia, and Africa. Substantial diversity among African isolates suggests this continent may be the region of origin of *B. abortus* as a species (JTF, unpubl. data). Inconsistencies in biovar assignment compared to their placement in the phylogenetic tree suggest that biovar classification in *B. abortus* does not consistently reflect genetic relationships or that biochemical determination of biovars is unreliable.

Six of our isolates from across Italy form a “Trans Italia” subclade along with isolates from Mongolia, southern Europe, and Egypt, which are defined by 44 SNPs. The broader clade also includes an isolate from Turkey. The 169 SNPs exclusive to this clade suggests a fairly long time period separates these isolates from the common ancestor they share with genomes from Uganda and Iraq. Isolates from eastern Italy form an ‘East Italia” subclade, with 157 SNPs exclusive to the branch. The broader clade contains an isolate from Greece, which shares 145 exclusive SNPs with the East Italia cluster. Members of the East Italia subclade are closely related, although these three isolates came from three different regions.

We developed 13 Melt-MAMA assays to genotype a larger collection of 261 samples and to better define the population substructure of *B. abortus* in Southern Italy. Six of the Melt-MAMA assays produced consistent results and corresponded to two assays each for the West Italia, Trans Italia, and East Italia subclades and the broader clades to which they belong (Table 1). Our genotyping indicated that 13 Italian isolates came from the West Italia subclade, 214 isolates were part of the Trans Italia subclade, and 34 isolates were part of the East Italia subclade (Supplementary material). We then mapped the geographic distributions of the clades to determine their phylogeographic patterns. The West Italia genotypes consisted entirely of isolates from biovar 1 detected in the three western Italian districts of Campania, Basilicata, and Calabria (Fig. 2). The East Italia group was found mostly in the eastern regions of southern Italy: Apulia, Molise and Abruzzo (Fig. 2). Finally, the Trans Italia group was distributed throughout endemic regions of Southern Italy, suggesting its dominance in the region (Fig. 2).

**Fig. 2.**
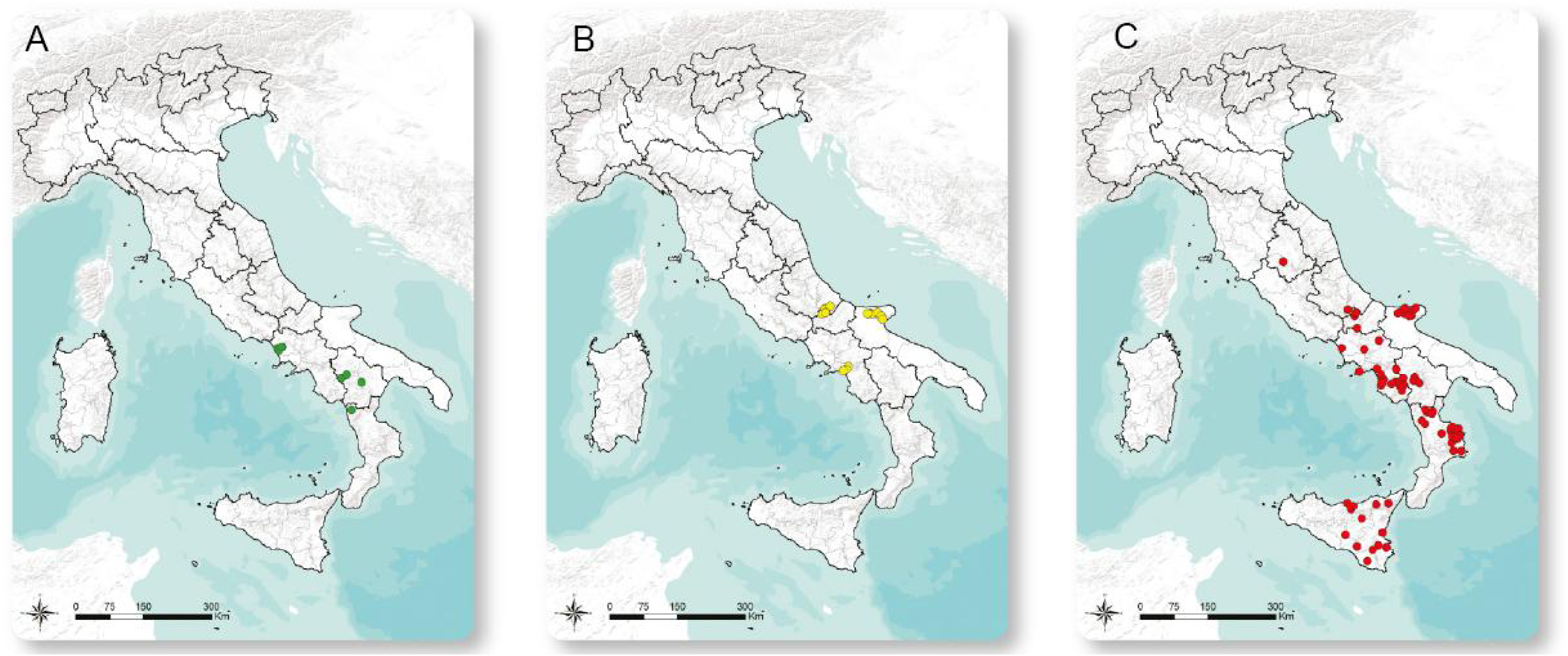
Geographical mapping of the Italian isolates according to Melt-MAMA genotyping. GIS coordinates separated by groups are shown as follow: green for the West Italia group (A), yellow for the East Italia (B) and red for the Trans Italia (C).

**Table 1.**
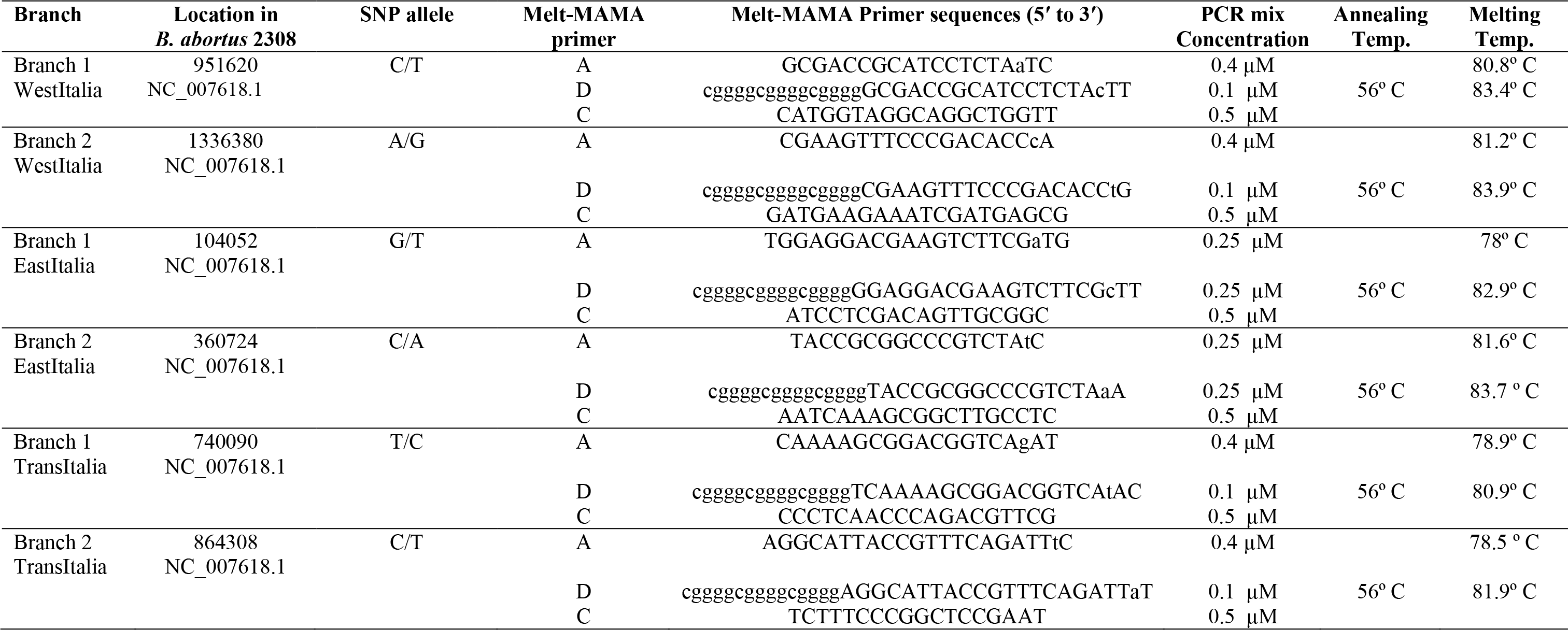
Characteristics of Melt-MAMA primers designed for six major branches containing Italian *Brucella abortus* isolates. Genome position is in the reference strain 2308 with chromosome number indicated in parentheses.

## Conclusion

Whole genome sequencing and comparison to a global collection of genomes allowed us to determine the evolutionary history of *B. abortus* in Italy. Despite the strains being isolated exclusively in 2011, the use of multi-locus variable number tandem repeat (VNTR) analysis (MLVA) in a previous study allowed us to select and then sequence a diverse collection of isolates from throughout endemic districts in Southern Italy [17]. Challenges with VNTRs limited our phylogenetic inference for *B. abortus* for the country. However, genome-wide phylogenetics using SNPs identified that Italian strains come from three major clades, all of which are substantially different than the common biovar 1/2/4 lineage found in much of Europe and North America. Genotyping assays supported these findings in a larger collection of 200 isolates from Italy.

We believe that the relatedness of Italian strains to lineages from other parts of Europe, Asia, and Africa suggests that *B. abortus* in Italy has a very different evolutionary history than the dominant lineage of biovar 1/2/4 that was brought to the New World by various Europeans in infected cattle. Two or three different domestication events of wild cattle in Eurasia have occurred [18], and then cattle, with their associated diseases, were introduced by humans throughout the world [19]. Italian farms contain several European cattle breeds, but also contain several primitive cattle breeds such as Podolica, Marchigiana, Chianina, and Romagnola, which are all directly linked to the ancient cattle *Bos primigenius* from Eurasia [20]. Current brucellosis programs strictly control cattle movements, but these programs have been only recently established, and *B. abortus* has likely been influenced by several centuries of introductions and disseminations of infected cattle throughout Italy.

The newly developed Melt-MAMA assays specific to the three Italian clades and subclades allowed us to screen a large collection of isolates and quickly and inexpensively assign them to one of three genetic groups. Future testing of additional isolates using these six SNP assays during eradication programs will allow them to be typed more rapidly and confidently than biotyping, and could serve Italian diagnostic laboratories as a first line assay. Genotyping coupled with knowledge of herd movement through use of risk maps, spatial modeling, and Social Network Analysis (SNA) techniques should enhance the reliability of the Italian brucellosis eradication plan. Although a multiplex PCR approach has been developed recently [14], and the MLVA-16 has been shown to be robust and informative [21], these techniques remain cumbersome for data production and standardization. Perhaps a more reasonable approach would be to pair the SNP assays with a limited number of VNTR loci to achieve a higher resolution for molecular epidemiology purposes. We recognize the capabilities of whole genome sequencing, but we are also aware that this technology is not yet accessible to all labs and requires skills that are not present in many diagnostic laboratories. Nonetheless, for some outbreak investigations, MLVA-16 or whole genome sequencing may be essential when complete epidemiological traceback is required. Ultimately, our study demonstrates the utility of

WGS SNPs paired with extensive epidemiologic data for analyzing the distribution of *B. abortus* isolates throughout endemic regions.

## Methods

### Aims of the study

The study was conducted to genetically characterize Italian *B. abortus* isolates from 261 animal cases in southern Italy from 2011 to 2014. The overall goal was to trace the origin and the global connections of bovine brucellosis in Italy and to develop a rapid genotyping method for novel isolates.

### *B. abortus* isolates

Large ruminants (cattle and water buffalo) and small ruminants (sheep and goat) testing positive for brucellosis in the serological tests carried out in the context of the national brucellosis eradication campaign are slaughtered, and the epidemiological investigation is supported through the isolation and characterization of *Brucella* spp. from selected tissues or body fluids. The preferred tissues for culture are lymphatic glands (i.e. mammary and genital lymph nodes, and spleen), uterus, and udder. Direct and enrichment *Brucella* cultures were carried out with these tissue samples following the OIE procedures. The suspected colonies were assigned to the species *B. abortus* and the relative biovar (1-7, 9) using PCR and traditional serotyping [22]. All cultured isolates were stored in the Italian collection with associated epidemiological data. To achieve maximum diversity necessary for these types of projects, we followed the guidelines Pearson et al. [23] to select 11 phylogenetically and geographically diverse *B. abortus* strains (Table 2) for whole genome sequencing using results from our VNTR study [17].

**Table 2.** Epidemiological data for 11 *Brucella abortus* isolates that were whole genome sequenced. All samples were collected in 2011 and were molecularly identified as *B. abortus*.

| **id** | **Region** | **Province** | **Town** | **Host** | **Biovar** |
| --- | --- | --- | --- | --- | --- |
| 1046 | Abruzzo | CH | Roccaspinalveti | cattle | 3 |
| 5586 | Calabria | KR | Pallagorio | cattle | 3 |
| 8486 | Campania | SA | Giffoni Valle Piana | cattle | 3 |
| 9060 | Calabria | CS | Scala Coeli | cattle | 3 |
| 8979-3 | Campania | CE | Cancello ed Arnone | water buffalo | 1 |
| 11796 | Sicilia | ME | Tusa | cattle | 3 |
| 12183 | Campania | SA | Corleto Monforte | cattle | 3 |
| 15074/1 | Puglia | FG | San Marco in Lamis | cattle | 3 |
| 1365/1 | Molise | IS | Carovilli | cattle | 3 |
| 3272 | Puglia | FG | Apricena | cattle | 3 |
| 9261 | Calabria | CS | Papasidero | cattle | 1 |

### Whole genome sequencing and SNP detection

We sequenced the isolates with an Illumina MiSeq to produce paired 300 bp reads with a 400 bp insert size (GenBank PRJNA284953) [24]. SNPs were determined with NASP version 1.0.0 [25], using default filters to remove SNPs from duplicated regions, read coverage less than 10X, base call proportion less than 90%, and loci not orthologous in all samples. NASP uses BWA [26] as the aligner and GATK [27] as the SNP caller. This analysis allowed us to compare these 11 genomes to 179 genomes of *B. abortus* from GenBank, accessed via PATRIC [28], that have been examined as part of a global analysis of *B. abortus* (JTF, unpubl. data). Phylogenetic trees were constructed using maximum parsimony in PAUP* 4.0b [29]. Branch support was assessed with 1000 bootstrap replicates and a consistency index was generated to assess the level of homoplasy.

### Melt-MAMA PCR assays

We designed SNP-specific assays that targeted six branches as shown in table 1 leading to specific Italian clades according to a maximum parsimony tree (Fig. 1). Downstream real time PCR analyses using Melt-MAMA were performed on 261 *B. abortus* isolates isolated in Italy from the endemic territories during the period 2011–2014 (Supplementary material). Briefly, Melt-MAMA allows sensitive genotyping assays to be developed that distinguish samples containing a single nucleotide difference, which can be robust determinants of specific lineages in clonal bacteria [30]. The assays satisfied the following five conditions as previously described [31], including: (i) Two 3´ forward primers containing sequences complementary to the allelic sequence for each SNP; (ii) a common reverse primer; (iii) an additional mismatch in the forward primer in anti-penultimate and penultimate position alternately; (iv) a GC-clamp in position 5´ for one of the forward primer; (v) amplicon of reduced size (60–120 bp). Relative primer concentrations were tested in ratios of 1:1, 4:1, and 1:4 during optimization of the PCR reaction conditions. Assays were considered valid when they gave consistently different melt profiles for samples containing SNPs for ancestral versus derived alleles.

## Declarations

### Ethics approval and consent to participate

This study is a retrospective molecular investigation of the *Brucella* collection from the National and OIE Reference Laboratory Istituto Zooprofilattico Sperimentale dell’Abruzzo e del Molise. No human or animal data were used therefore informed consent was not required.

### Consent for publication

Not applicable.

### Availability of data and materials

The genomes are available at GenBank under the project PRJNA284953

### Authors’ contributions

GG and JTF have designed the study and drafted the final manuscript, EDG, FDM, CC and PC contributed in planning the project and critically reviewed the manuscript, IP, AA, LS, KZ,GDD and MA performed the WGS, Melt-MAMA and conventional tests for typing the *Brucella* isolates, GG, JTF and KPD conducted the bioinformatics analyses. All authors read and approved the final manuscript.

## Acknowledgments

We thank Istituti Zooprofilattici Sperimentali (IIZZSS) for the technical support. We are grateful to Paola Di Giuseppe for providing assistance on the graphics.

## List of abbreviations

PCR.: Polymerase chain reaction;
VNTR.: Variable number tandem repeat;
MLVA.: Multi Locus VNTRs Analysis;
BSL.: biosafety level 3 laboratory;
SNPs.: single nucleotide polymorphisms.
Melt-MAMA.: Melt analysis of mismatch amplification mutation assays.

